# Expanding plant trait databases using large language model: A case study on flower color extraction

**DOI:** 10.1101/2025.02.11.637746

**Authors:** Masaru Bamba, Shusei Sato

## Abstract

Plant trait databases play a crucial role in understanding ecological and evolutionary processes yet remain insufficient due to geographical and data availability limitations. To address this limitation, we developed a novel large-scale text extraction approach using a large language model (LLM) to transform descriptions of Floras into structured trait data, thereby expanding existing databases.

We applied this approach to extract flower color information from the *Flora of China* and integrate it into the TRY Plant Trait Database. After integration, the dataset expanded to 27,252 species, more than doubling the previously available flower color records. Additionally, we linked the dataset with occurrence records from GBIF and environmental data, including climate and soil properties, to disentangle ecological insight and flower color distribution.

Our large-scale association analysis of flower colors and environments revealed that white, yellow, and red-type flowers exhibit distinct environments, suggesting that abiotic environment can play a role in flower color evolution.

By transforming descriptions of Flora into structured data, our approach organizes traits across more plant species, creating new opportunities for ecological and evolutionary research. The present approach can be extended to other traits, enhancing our understanding of how plants adapt and respond to environmental changes on a global scale.

## Introduction

In recent years, climate change and environmental modification have progressed on a global scale, leading to increasing expectations for research that integrates large volumes of biological and environmental data to better understand ecosystems and landscapes, uncover mechanisms governing the establishment and maintenance of biological communities, and clarify evolutionary processes in various species (Hampton et al., 2013). Consequently, extensive efforts to collect diverse biological information are in progress. For example, the Global Biodiversity Information Facility (GBIF; http://www.gbif.org) makes available approximately five million species and three billion specimen/observation records. In the field of plant science, the TRY Plant trait database project (https://www.try-db.org) was launched in 2007, and has assembled 686 datasets containing phenotypic information for 305,594 plant taxa and 15,409,681 traits. Both these databases provide valuable information and continue to expand their datasets.

In this study, we propose extracting information from plant Floras—specialized publications that systematically document and describe the plant species of a particular region— as one approach to expanding plant-related datasets. Flora is a high-quality, expert-reviewed resource that organizes scientific names alongside morphological characteristics and habitat information, all of which are valuable for research. However, most floras are written in natural language, which complicates handling. The recent growth of Large Language Models (LLMs) has made it possible to automatically extract meaningful knowledge from textual sources and convert it into structured database entries (Dagdelen et al., 2022). Building on these advances, our goal is to use LLMs to efficiently process extensive flora information, thereby constructing a large-scale dataset that can complement existing databases.

We focus on Flora of China (Brach and Song, 2006; http://www.efloras.org) to address gaps in current plant databases. Although the TRY database has improved since its launch, it still tends to be biased toward species distributed in Europe and the Americas (Kattge et al., 2022). Consequently, adding more information on Asian flora, which is notably diverse, holds considerable value. Indeed, East Asia exhibits a high level of plant diversity (Qian and Ricklefs, 2000), and around 31,000 species distributed at China alone, one-eighth of the world’s total (Brach and Song, 2006). Compiled as an international collaborative project, Flora of China aims to provide comprehensive coverage of China’s diverse flora and offers expert-verified phenotypic information. Here, we extract this body of text using an LLM to construct a large-scale dataset that can enhance existing resources.

To assess the advantages of the constructed dataset, we examine the relationship between flower color and environmental factors. Flower color is one of the most visually prominent traits in angiosperms and frequently appears in species descriptions because it shows considerable diversification even among closely related plants (Rausher, 2008). Therefore, it provides an ideal model for examining approaches to extracting information from textual sources. Moreover, although flower color is considered ecologically important, its evolutionary processes are not fully understood, indicating its research significance. While flower color has long been regarded as a trait predominantly shaped by pollinator-mediated selection, the pigments responsible for coloration are also known to accumulate in response to various environmental stresses. It is therefore likely that flower color exposed multidirectional selection pressures beyond pollinator preferences (Rausher and Fry, 1993; Simms and Bucher, 1996; Fineblum and Rausher, 1997; Armbruster, 2002; Irwin et al., 2003; Strauss and Whittall, 2006). Recent studies have even reported that flower color is not always a reliable indicator of pollination syndromes (Dellinger, 2020), suggesting that factors unrelated to pollination may contribute to its evolution. However, there have been few large-scale studies examining the relationships between flower color and environmental factors, and it remains unclear whether differences in flower color correlate with distinct habitat environments.

Here, we integrated flower color data from the TRY database with flower color information extracted from Flora of China via ChatGPT (OpenAI, 2024). We then analyzed species occurrence data obtained from GBIF, along with climate information from WorldClim (Fick and Hijmans, 2017) and soil data from ISRIC SoilGrids (Poggio et al., 2021). By synthesizing this information in a statistical model, we evaluated the relationship between flower color and environmental factors on a large geographical scale.

## Materials and Methods

### Collection of Flower Color and Habitat Information

The flower color information used in this study was obtained by integrating (1) data from the TRY database and (2) information extracted from the Flora of China (Fig. 1). From the TRY database, we acquired data corresponding to TraitID = 207 (TraitName = “Flower color”), retaining the contents of the column “OrigValStr” as the flower color information. The six datasets that included flower color information were: the BiolFlor Database, PLANTS data USDA, Orchid Trait Dataset, Trait Data for African Plants – a Photo Guide, Functional Flowering Plant Traits, and Traits of Urban species from Ibagué Columbia.

**Figure 1.**
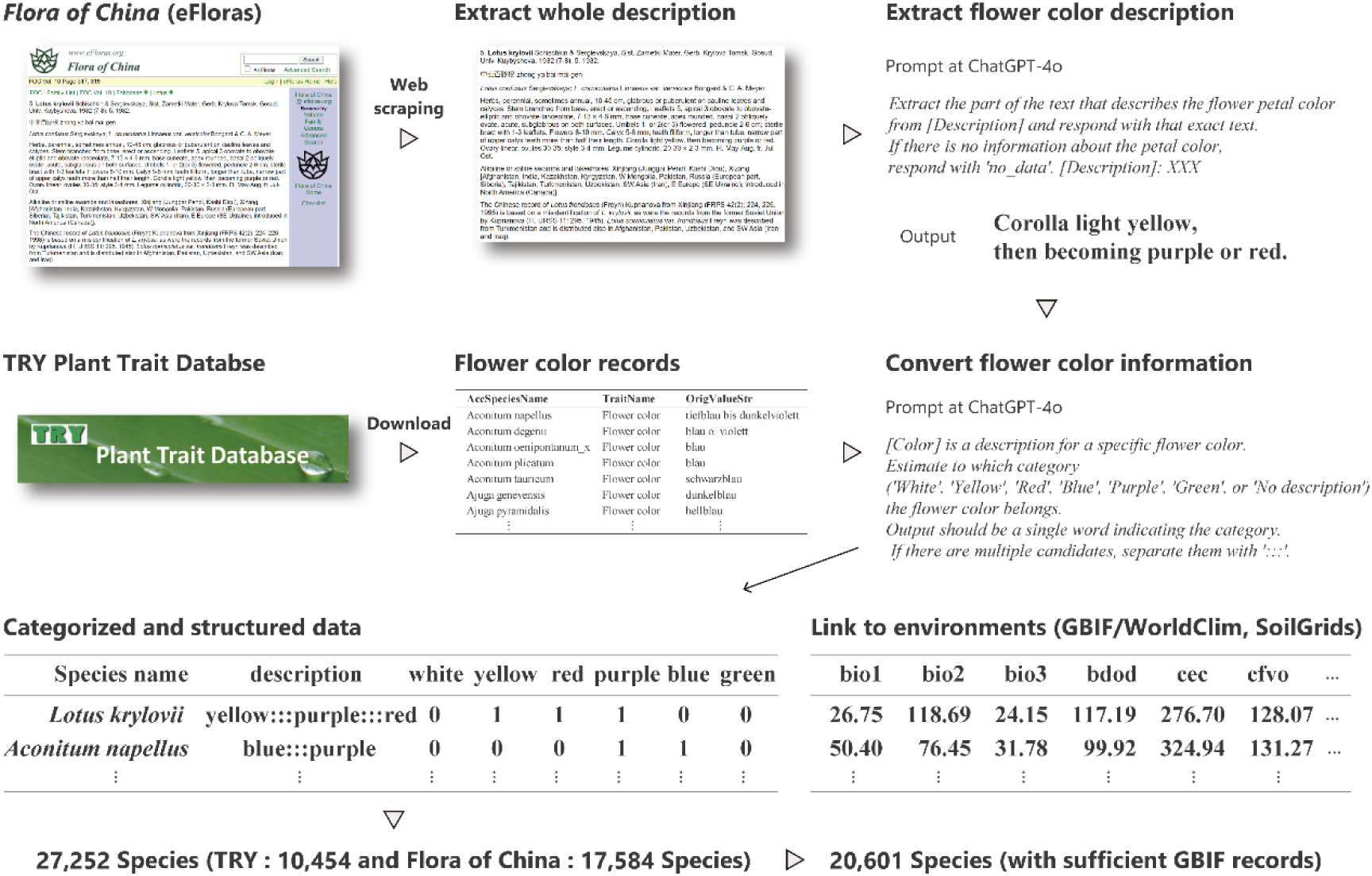
Overview of the data extraction and integration workflow.

To collect flower color information from the Flora of China, we first performed web scraping of all taxonomic descriptions published in Volumes 4–25 (excluding Volume 1, which is introductory, and Volumes 2–3, which deal with Pteridophytes) and obtained the descriptive text (https://github.com/mbamba2093/Flower-Color-Habitat-Environment-Association/01_Scraping_FloraOfChina.py). Next, among the retrieved text, only the sections related to flower color were extracted using ChatGPT-4o. The prompt used was as follows (where XXX represents the previously obtained descriptive text):

> *“Extract the part of the text that describes the flower petal color from [Description] and respond with that exact text. If there is no information about the petal color, respond with ‘no_data’. [Description]: XXX”*

Although the prompt explicitly mentions “flower petal color,” we confirmed that descriptions pertaining to other floral components (such as the corolla) were also captured. For further details, refer to the GitHub repository at https://github.com/mbamba2093/Flower-Color-Habitat-Environment-Association/02_Extract_flower_color_description.py.

We then converted the flower color information obtained from the TRY database and the Flora of China into predefined categories using ChatGPT-4o. The prompt used was as follows:

> “*[Color] is a description for a specific flower color. Estimate to which category (’White’, ‘Yellow’, ‘Red’, ‘Blue’, ‘Purple’, ‘Green’, or ‘No description’) the flower color belongs. Output should be a single word indicating the category. If there are multiple candidates, separate them with ‘:::’.*”

If there were inconsistencies in the category labels output by the model, they were manually corrected. Furthermore, if the category included any of “Red,” “Blue,” or “Purple,” it was grouped into “RedType.” In subsequent analyses, we focused on whether each plant species possessed any of the following flower color elements: “White,” “Yellow,” or “RedType.” The scripts used and the manual conversion process are compiled at https://github.com/mbamba2093/Flower-Color-Habitat-Environment-Association/03_Categorize_Flower_color.py. Additionally, for each plant species with flower color information, we used the *taxize* package (Chamberlain and Szocs, 2013) in R to append taxonomic information and verified that the species were angiosperms.

To assess the consistency between the flower color data contained in the TRY database and those extracted from the Flora of China, we evaluated two metrics for plant species that were observed in both datasets. The first metric was whether the category information (“White,” “Yellow,” or “RedType”) was a complete match (“Complete correct”). The second was the percentage agreement with respect to the three categories (“Correct rate”). These metrics were applied not only to comparisons between the Flora of China and each dataset in the TRY database, but also to comparisons among different datasets within the TRY database. Moreover, we compared the integrated (union) set of flower color information from the entire TRY database with that of the Flora of China. In the subsequent analyses, we used the union of the flower color data from the TRY database and the Flora of China.

For plant species whose flower color information was successfully obtained, we retrieved habitat data from the GBIF database using *rgbif* (Chamberlain, 2019). With *rgbif*, we gathered up to 10,000 occurrence records (latitude/longitude). We excluded data flagged by the GeospatialIssue indicator as defined by GBIF. From the occurrence data obtained via GBIF, we extracted and combined the following environmental information based on each occurrence’s latitude and longitude:

1. **Climate information**: Bio1–Bio19 and elevation data from WorldClim (10 km resolution; Fick and Hijmans, 2017).
2. **Soil information**: From SoilGrids (250 m resolution; Poggio et al., 2022), we obtained the following variables for the layer closest to the surface: bdod (bulk density of the fine earth fraction), cec (cation exchange capacity), nitrogen (total nitrogen), phh2o (soil pH), soc (soil organic carbon content), ocd (organic carbon density), ocs (organic carbon stock), clay (clay fraction), silt (silt fraction), sand (sand fraction), and cfvo (coarse fragment volume).

For specimens that came from the same climate and soil grid cells, we removed duplicates. We included plant species with six or more specimens or observation records in GBIF in our subsequent analyses, assuming sufficient data availability on GBIF for these species.

As representative values of each plant’s habitat environment, we calculated a 20% trimmed mean for each environmental variable. For the soil environment, we computed pairwise similarities (Euclidean distances) among the soil variables and took the average distance as an indicator of how diverse the soil environments in which the species occurs. Next, we carried out a PCA on all species’ climate, soil, and elevation, using scores of the first to tenth principal components (PC1–PC10) as our environmental axes.

For species with flower color and habitat information, phylogenetic data were obtained from TimeTree 5 (Kumar et al., 2022). Synonyms and taxa with insufficient information were replaced by appropriate taxa according to TimeTree standards. Any branches in the phylogeny that had a length of zero were replaced with the smallest non-zero branch length, after which the branch lengths were recalculated using Grafen’s method, resulting in an ultra-metric tree for subsequent analyses.

### Statistical Analyses

To explore the relationship between environmental axes and flower color elements, we employed a hierarchical threshold model implemented via *MCMCglmm* (Hadfield 2010). As the response variable, we used the presence or absence of each flower color element (“White,” “Yellow,” or “RedType”) encoded as binary (0/1). As explanatory variables, we incorporated the aforementioned environmental axes (PC1–PC10). Based on the ultra-metric tree we created, we computed the inverse covariance matrix and included it as a random effect in the model. For the prior distribution, we set the initial variance of the random effect to 1 and the degrees of freedom of the inverse-Wishart distribution to 0.002. We ran 1,050,000 MCMC iterations, discarding the first 50,000 as burn-in and sampling every 100 iterations. Convergence was confirmed using Gelman– Rubin diagnostics and Raftery–Lewis diagnostics. We also conducted similar *MCMCglmm* analyses, each time using only one environmental variable as an explanatory variable, to further clarify individual environmental factors associated with flower color elements.

To account for potential phylogenetic confounding, we subdivided the data by plant family and performed the same *MCMCglmm* analyses within each family that had 50 or more species. We also applied a GLM without random effects to the entire dataset to compare with results that ignore phylogenetic information. Furthermore, for datasets including species for which it was difficult to obtain phylogenetic information, we constructed a similar model treating family and genus as random effects in *MCMCglmm* and also explored a GLM without random effects. In doing so, we aimed to minimize potential bias that could arise from excluding species lacking phylogenetic data.

The R code used for the above analyses is available on the following GitHub repository: https://github.com/mbamba2093/Flower-Color-Habitat-Environment-Association/.

## Results

In this study, we accessed descriptive information for 37,723 species in the Flora of China and obtained flower color data for 17,584 of these species. The TRY database contained flower color entries from six datasets, providing 20,434 records in total (5,890 from the BiolFlor Database, 2,102 from PLANTS data USDA, 17 from the Orchid Trait Dataset, 12,068 from Trait Data for African Plants – a Photo Guide, 155 from Functional Flowering Plant Traits, and 202 from Traits of Urban species from Ibagué Columbia), covering 10,454 species. By combining these, we compiled flower color information for 27,252 species (321 families, 3,293 genera; Table S1). Of these, 20,601 species had sufficient habitat information from GBIF (Fig. 2), and phylogenetic information was available for 7,938 species.

**Figure 2.**
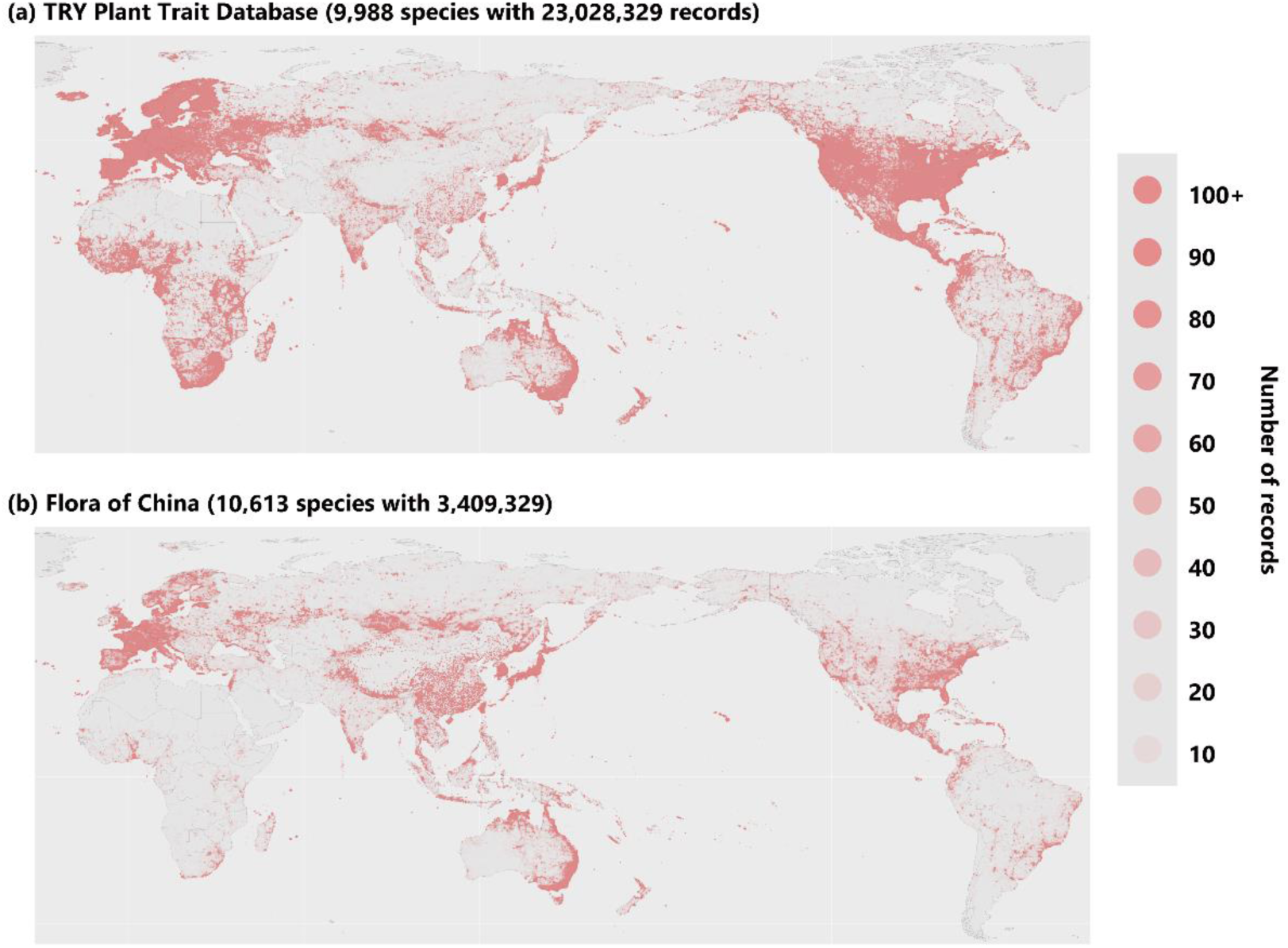
Distribution of occurrence records. Red circles indicate the locations where GBIF records have been reported for the plant species used in this study. (a) Occurrence records for 9,988 species with flower color information recorded in the TRY Plant Trait Database. (b) Occurrence records for 10,613 species for which flower color information was extracted from Flora of China in this study.

Each plant species was classified based on whether its flowers possessed any of the following color elements: white, yellow, red, blue, purple, or green. White was the most common (8,710 species), followed by yellow (7,414 species) and red (5,103 species) (Fig. 3A). For the analyses, flowers with any element of red, blue, or purple were grouped as “RedType,” resulting in 9,689 species. Conditional probability calculations indicated that white elements frequently co-occur with RedType (P(White|RedType) = 0.7370, P(RedType|White) = 0.7289), whereas yellow tended to co-occur less with other colors (P(Yellow|White) = 0.4251, P(Yellow|RedType) = 0.4187, P(White|Yellow) = 0.6204, P(RedType|Yellow) = 0.6043) (Fig. 3B).

**Figure 3.**
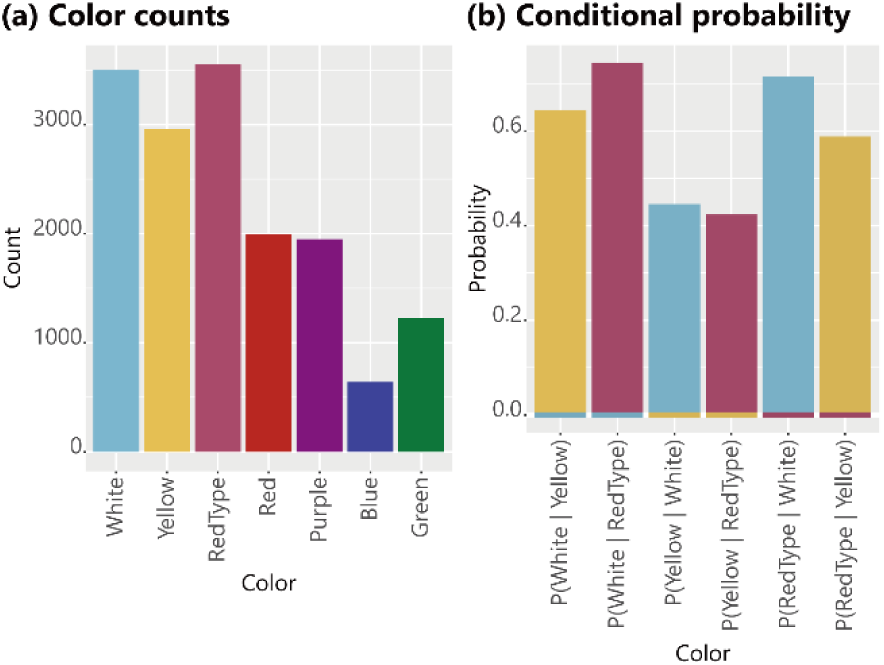
Counts of flower color data. (a) Counts of flower color records used in this study. (b) Conditional probabilities between individual flower colors in cases where multiple colors were present.

A total of 773 species had overlapping records in *Flora of China* and the TRY database. Among these, 55.7% (431 species) showed complete agreement in flower color elements (Table 1), and the match rate across the three categories (White, Yellow, RedType) was 82%. Moreover, examining three datasets within the TRY database (BiolFlor, PLANTS data USDA, and Trait Data for African Plants) revealed complete match rates of 41% to 61% and category match rates of 71% to 83%. These observations suggest that differences between *Flora of China* and the TRY database are comparable to the variability among TRY datasets themselves.

**Table 1.** Correct rates among datasets.

| Dataset 1 | Dataset 2 | Overlap<br>species | Consistent<br>species | Average element-wise<br>matching |
| --- | --- | --- | --- | --- |
| Flora of China | TRY all database | 773 | 431 | 0.8202 |
| Flora of China | BiolFlor Database | 405 | 250 | 0.8494 |
| Flora of China | PLANTS data USDA | 166 | 82 | 0.7550 |
| Flora of China | Functional Flowering Plant Traits | 44 | 26 | 0.8258 |
| Flora of China | Traits of urban species from Ibagué<br>Colombia | 36 | 18 | 0.7593 |
| Flora of China | Trait Data for African Plants | 317 | 160 | 0.8013 |
| BiolFlor Database | PLANTS data USDA | 184 | 110 | 0.8098 |
| Trait Data for African<br>Plants | PLANTS data USDA | 75 | 31 | 0.7156 |
| Trait Data for African<br>Plants | BiolFlor Database | 57 | 35 | 0.8363 |

Next, we investigated the relationship between flower color and habitat environment using 7,938 species (307 families, 2,231 genera) for which phylogenetic and environmental data were available. We employed a Bayesian linear mixed model with *MCMCglmm*, treating flower color as the response variable, compressed environmental data (PC scores) as the explanatory variables, and phylogenetic covariances based on the phylogeny as a random effect. Gelman–Rubin diagnostics indicated near-unity values for all parameters (Multivariate potential scale reduction factor [MPSRF] for White: 1.00051, Yellow: 1.000729, RedType: 1.000982), and trace plots were stable (Fig. S1), suggesting that the model converged successfully. In addition, the random-effect variance exceeded 1 for all flower colors (95% confidence intervals of the variance–covariance matrices were White: 119.8969–120.6105, Yellow: 217.0301– 218.2140, RedType: 187.2230–188.2368), indicating the statistical importance of phylogenetic relationships for flower color.

The *MCMCglmm* results revealed distinct relationships between each flower color and certain environmental PC axes (Fig. 4A, Table S2). For white flowers, PC1, PC2, and PC5 showed positive effects, while PC7 was negative. For yellow flowers, PC2, PC4, PC6, PC7, and PC9 had negative effects. In the case of RedType flowers, PC1, PC2, PC4, PC6, and PC7 had negative effects, whereas PC3 was positive. Notably, PCs 1 to 3 differed among the colors, indicating that different flower colors distributed in different environments.

**Figure 4.**
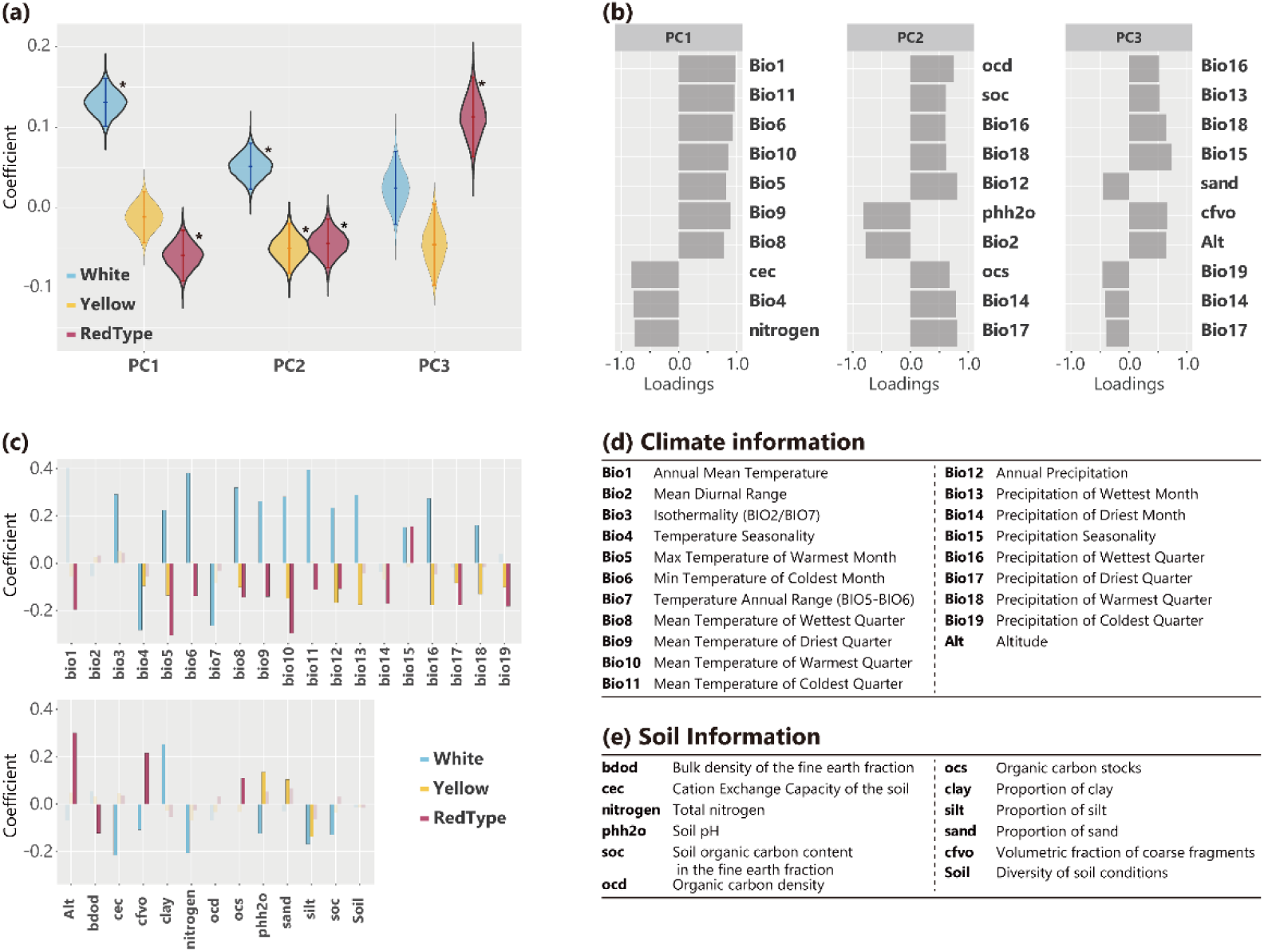
Coefficients of environmental factors for flower colors. (a) The vertical axis represents the coefficients obtained from the MCMCglmm analysis, showing only the fixed effects for PC1–PC3. The violin plots illustrate the posterior distribution of the coefficients sampled during the MCMC process. The bars within the violin plots indicate the 95% credible intervals, while the points represent the mean values. Violin plots outlined with a thick line and accompanied by an asterisk denote significant coefficients, where the 95% credible interval does not overlap with zero. The violin plots are categorized into three groups corresponding to white, yellow, and red-type flowers. (b) Factor loadings of environmental variables on the PCA axes. The plot shows the top 10 variables with the highest absolute factor loadings for PC1– PC3, which were used as environmental axes in the model. (c) Estimated coefficients for individual environmental variables. The bar plot displays the estimated coefficients for each environmental variable when analyzed separately in the model. For each variable, coefficients are shown for white, yellow, and red-type flowers. Bars outlined with a border indicate significant coefficients, where the 95% credible interval does not overlap with zero. (d) and (e) Abbreviations of environmental variables used in the analysis. (d) Climate variables obtained from WorldClim. (e) Soil variables obtained from SoilGrids.

Summarizing factor loadings for these PC axes and their relationships with specific environmental factors revealed distinct environmental conditions for each flower color. Figure 4B summarizes the factor loadings for PC1 through PC3. In total, 22 significant variables were identified for white flowers, 13 for yellow flowers, and 16 for RedType flowers (Fig. 4C, Table S3). The following environmental factors appeared in the top 10 loadings for PC1 through PC3 and were significant in single variable models for each flower color (Fig. 4D, E):

- White (Positive: Bio5, 6, 8, 9, 10, 11, 12, 16, 18; Negative: Bio4, cec, nitrogen, phh2o)
- Yellow (Positive: phh2o; Negative: Bio12, 16, 17, 18)
- RedType (Positive: Alt, cfvo; Negative: Bio1, 5, 6, 8, 9, 10, 11, 12, 14, 15, 17, 19)

To test the robustness of these findings against phylogenetic confounding, we conducted *MCMCglmm* analyses for individual families and also performed GLM analyses without random effects. In the family-level *MCMCglmm* models, the MCMC converged in six families for white flowers (Asteraceae, Campanulaceae, Ericaceae, Fabaceae, Orchidaceae, and Rosaceae), five for yellow flowers (Apiaceae, Fabaceae, Orchidaceae, Poaceae, and Rosaceae), and seven for redtype flowers (Apiaceae, Asteraceae, Ericaceae, Fabaceae, Orchidaceae, Poaceae, and Rosaceae) (Table S3). Not all PC1–PC3 axes were significant in every combination, but the results were not in conflict with the full-dataset analysis (Table S4). The GLM results aligned with those of *MCMCglmm* (Table S5, and the effect sizes for PC1–PC3 ranged in the *MCMCglmm* confidence intervals, with the exception of PC1 for white flowers. These observations confirm that the influence of environmental axes on flower color is robust to phylogenetic confounding.

We performed additional analyses for 20,601 species that included those lacking available phylogenetic information. In an *MCMCglmm* model that treated family and genus as random effects, white flowers showed positive effects for PC1 and PC2, yellow flowers showed a negative effect for PC2, and red flowers showed negative effects for PC1 and PC2 and a positive effect for PC3 (Table S6)—results that were consistent with the analysis with phylogenetic information. Likewise, GLM models without random effects yielded no contradictions, and the effects of PC1–PC3 for flower color were within the corresponding confidence intervals (except PC1 for white flowers and PC3 for red flowers; Table S7). These findings suggest that the results do not arise solely from the subset of species with available phylogenetic information.

## Discussion

The dataset produced by this research provides a global resource for analyzing flower color and habitat in plants. Our work improved the geographical representation in existing databases by integrating additional material from Asia, reducing the emphasis on Europe and North America. Specifically, we developed an automated pipeline that uses an LLM to extract color information from the Flora of China description, resulting in a large-scale dataset covering 27,252 species. Of these, 20,601 include environmental details, allowing for comprehensive analyses of relationships between flower color and environment. This dataset provides a valuable basis for advancing research on the ecological and evolutionary roles of flower color and its interactions with the environment.

The LLM-powered pipeline we developed can be applied to other Floras and phenotypic traits beyond flower color research. Flora, field guides, and other specialized publications contain expert-validated knowledge, but the process of transforming such natural language information into structured data has traditionally been time-consuming and resource-intensive. By applying an LLM’s ability to accurately interpret text, we were able to transform natural-language descriptions into structured datasets. Moreover, LLMs can process multiple languages, making them useful for handling multilingual databases. For example, the TRY database used in this study contains data in English, German, and Spanish, which can be managed more efficiently with LLMs. LLMs are also useful for assigning phenotypic information to categories, making it easier to handle complex biological datasets. This approach can be extended to other traits such as leaf shape, seed size, or growth forms, broadening the scope of future research.

Our large-scale dataset reveals significant relationships between flower color and the environment, with distinct patterns emerging for each flower color. Statistical modeling and PCA factor loadings suggest that white flowers are more common in locations with higher temperatures, greater precipitation, lower soil pH, greater soil carbon content, and lower nitrogen and cation exchange capacity. Yellow flowers tend to appear in areas with less precipitation, lower soil carbon, and greater daily temperature fluctuations. Additionally, RedType flowers are more frequent in cooler habitats with lower soil carbon, higher nitrogen and cation exchange capacity, higher elevations, and greater seasonal variability in rainfall. These findings imply that different lineages may have spread into distinct environments according to their respective flower colors. In addition, results obtained at the family level did not contradict the overall findings, and the effects were robust to phylogenetic confounding, indicating that these color–environment relationships are prevalent across plant taxa.

The relationships between flower color and environment may be explained by the properties of flower pigments. For instance, anthocyanins (common in red, blue, and purple flowers) and carotenoids (common in yellow flowers) can accumulate in response to environmental stresses due to their antioxidant properties. Anthocyanins are known to increase in various plant organs under drought conditions (Warren and Mackenzie 2001; Strauss and Whittall 2006; Warren and Mackenzie 2001), low temperatures (Christie et al. 1994; Pietrini et al. 2002; Stiles et al. 2007; Liu et al. 2018), and high ultraviolet exposure (Mtileni et al. 2024), and carotenoids also accumulate in dry conditions (Munné-Bosch and Alegre 2000). Therefore, in areas with limited rainfall, flower colors with anthocyanins and carotenoids may be selected. Furthermore, because anthocyanin synthesis and degradation can occur rapidly (Hughes et al. 2005), it could serve an adaptive role in environments with highly variable conditions. White flowers, meanwhile, may reflect more light (Stickland 1974) and have lower production costs (Carlson and Holsinger 2010, 2013). This could help regulate temperature around reproductive organs in warm, humid areas where evaporation is restricted, and it may also benefit plants growing in soils that are deficient in nitrogen and cation exchange capacity.

Although this study identifies general patterns in the relationship between flower color and the environment, some cases deviate from these trends. For instance, anthocyanin accumulation can increase at high temperatures (Dela et al. 2003), and white flowers may dominate at high elevations (Mogford 1974). It remains unclear whether flower color distributions primarily result from direct selection on flower color itself or from genetic traits influencing pigment deposition across various plant tissues. While natural selection may act directly on flower color, it is also possible that selection favors the accumulation of specific pigments in multiple plant parts. In such cases, acquiring genetic predispositions that facilitate pigment accumulation could play a crucial role in shaping flower color evolution. Further research, including the accumulation of additional cases across various species, is needed to clarify these mechanisms and determine the conditions under which these general patterns prevail or whether other selective pressures become dominant.

Finally, although this research integrated extensive data, some limitations remain. For instance, our current dataset cannot determine whether the observed relationships are driven by direct environmental influences or by pollinators favoring certain flower colors and environments, as it does not include pollinator information. Moreover, coverage of the Oceania region and endemic species in certain locations remains limited, making it difficult to draw fully global conclusions. Going forward, gathering data from these underrepresented regions and incorporating pollinator-related details could offer a more comprehensive understanding of how flower color evolves in response to environmental pressures. Such advances would be invaluable for both evolutionary ecology and conservation biology, enhancing our understanding of plant-environment interactions.

## Acknowledgments and Funding

This work was supported by JSPS KAKENHI [grant number 21K14763; 24K18168 to MB, JP20H2884 to SS]

## Author Contributions

MB planned and designed the research. MB performed research. MB and SS wrote the manuscript.

## Data Availability Statement

All data, program code, figures, tables, and supplemental information used in this study are available from the GitHub repository. https://github.com/mbamba2093/Flower-Color-Habitat-Environment-Association

## Conflict of Interest

None declared.

## Figure legends

**Figure S1.**
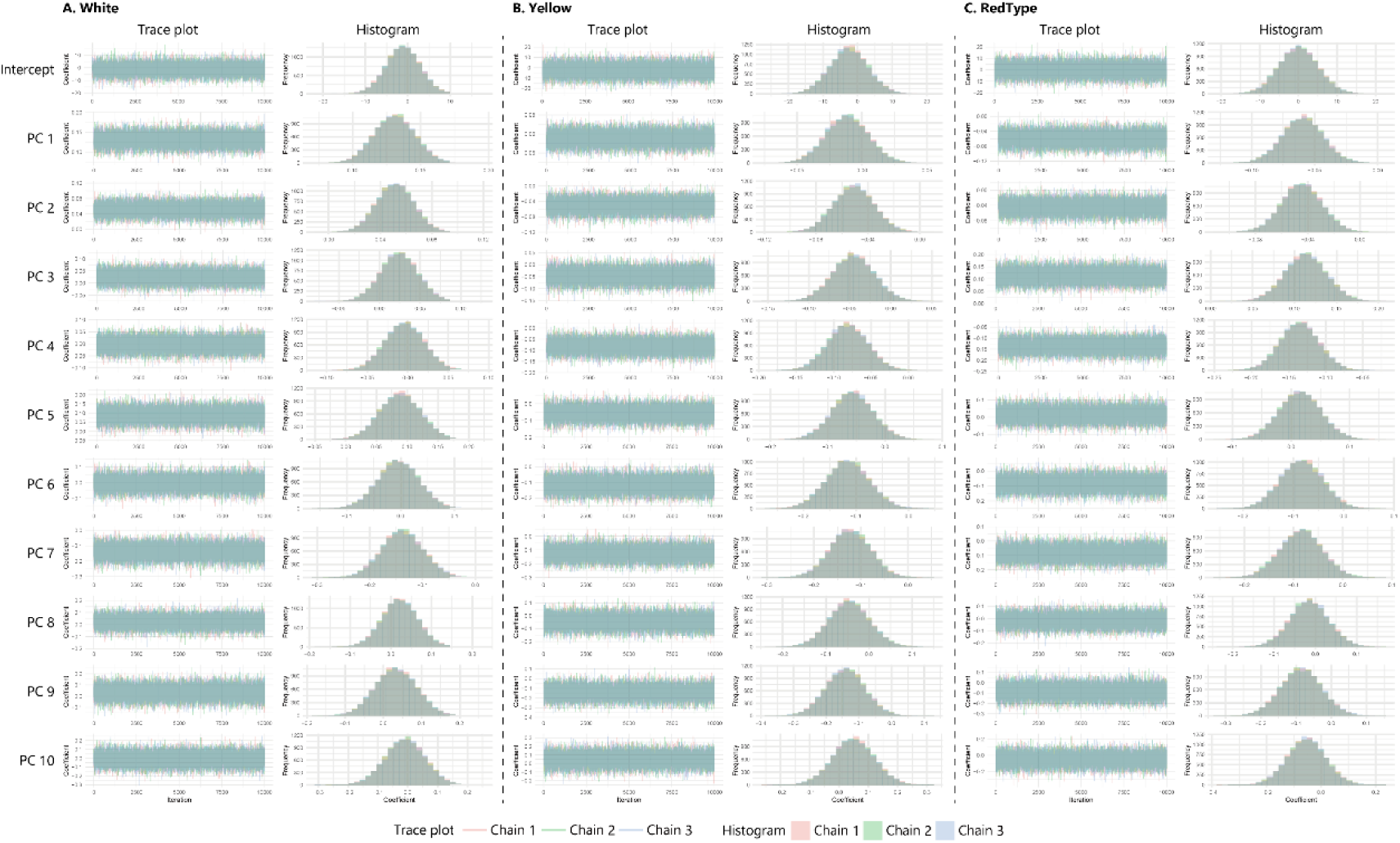
Trace plots from the MCMCglmm analysis. Trace plots and posterior distributions for each flower color and variable are shown. Different colors represent Chain 1–3.

## Notes

https://github.com/mbamba2093/Flower-Color-Habitat-Environment-Association

### Competing Interest Statement

The authors have declared no competing interest.

